# Uncovering the Mechanism of the Proton-Coupled Fluoride Transport in the CLC^F^ Antiporter

**DOI:** 10.1101/2022.09.27.509787

**Authors:** Kira R. Mills, Hedieh Torabifard

## Abstract

Fluoride is a natural antibiotic abundantly present in the environment and, in micromolar concentrations, is able to inhibit enzymes necessary for bacteria to survive. However, as is the case with many antibiotics, bacteria have evolved resistance methods, including through the use of recently discovered membrane proteins. One such protein is the CLC^F^ F^-^/H^+^ antiporter protein, a member of the CLC superfamily of anion-transport proteins, most notably known for their ability to transport chloride ions. While it possesses many similarities to the other CLC proteins, it also differs in several key ways, and though previous studies have examined this F^-^ transporter, many questions are still left unanswered. To reveal details of the transport mechanism used by CLC^F^, we have employed molecular dynamics simulations and umbrella sampling calculations. Our results have led to several discoveries, including the mechanism of proton import and how it is able to aid in the fluoride export. Additionally, we have determined the role of the previously identified residues Glu118, Glu318, Met79, and Tyr396. This work is among the first computational studies of the CLC^F^ F^-^/H^+^ antiporter and is the first to propose a mechanism which couples both the proton and anion transport.

## Introduction

Fluoride is well-known to act as a natural antibiotic, inhibiting the enolase and phosphoryl-transfer enzymes necessary for bacterial survival, at concentrations as low as 10-100 μM.^1^ It’s unsurprising, then, that researchers recently discovered a riboswitch which, upon F^-^ binding, turns on transcription for two phylogenetically unrelated bacterial fluoride exporters: the CLC^F^ F^-^/H^+^ antiporter, belonging to the CLC superfamily of anion-transport proteins; and Fluc, a fluoride-specific ion channel.^2,3^ Since their discovery in 2012, there have been few studies examining these proteins,^1,2,4–12^ leaving many questions unanswered.

Previous studies on the CLC^F^ protein^4,6,7^ have shown that despite having many similarities to the well-studied CLC Cl^-^/H^+^ antiporter, it is still significantly different beyond just the selectivity for F^-^ over Cl^-^ and a unique stoichiometry, exchanging one F^-^ for each H^+^ instead of the canonical 2:1 anion to proton ratio.^2,13,14^ Figures 1A and 1B highlight the CLC Cl^-^/H^+^ antiporter binding area, and its mechanism has been shown to involve a gating external glutamate (Glu148) which gates the anion path, while also transferring the transported H^+^ to an internal glutamate (Glu203), in a well-described bifurcated pathway.^15^ It’s been shown that Glu148 of the CLCs is able to rotate clockwise and pick up a proton before donating it to Glu203 via a transient water wire.^16^ It can then continue rotating, and its negative charge is able to push the Cl^-^ anions out of the protein.^17^ The transport pathway has also been shown to possess three distinct anion binding sites (S_ext_, S_cen_, and S_int_) with important contributions coming from conserved serine (Ser107) and tyrosine (Tyr445) residues.^18^

**Figure 1:**
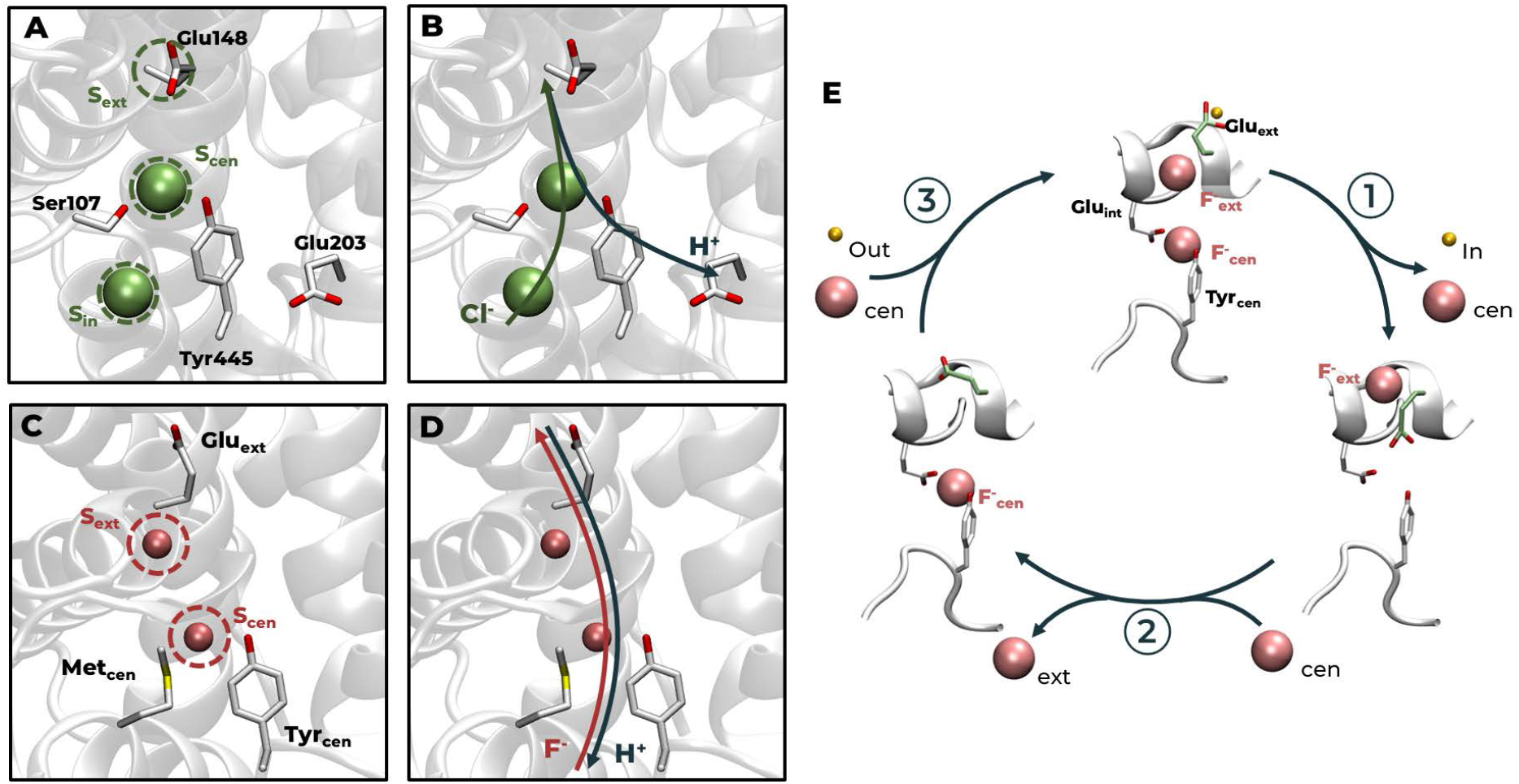
A. Binding sites and important residues in CLC Cl^-^/H^+^ antiporter, binding sites are circled in green; B. Bifurcated transport pathway in CLC Cl^-^/H^+^ antiporter; C. Binding sites and important residues in CLC^F^ F^-^/H^+^ antiporter, binding sites are circled in pink; D. Previously proposed single transport pathway in CLC^F^ F^-^/H^+^ antiporter; E. Windmill model proposed by Last *et al*.^6^ with Glu_int_ and Tyr_cen_ shown in white licorice and Glu_ext_ in green licorice to highlight its rotation. Fluoride ions are represented as pink spheres, chloride ions are represented as green spheres, and the proton is represented as a yellow sphere.

In the CLC^F^ protein, however, initial studies show that while it possesses an external glutamate (E118, hereafter referred to as Glu_ext_) and a tyrosine (Y396, hereafter referred to as Tyrcen) in the same positions, it is lacking both the internal glutamate and the conserved serine, and instead possesses a methionine (M79, hereafter referred to as Met_cen_) in the approximate serine location.^7^ Additionally, only two anion binding sites have been determined, S_ext_ and S_cen_,^6^ shown in Figure 1C.

These details combined with mutant studies have given way to a proposed windmill transport model which indicates just a single pathway,^6^ depicted in Figures 1D and 1E. Specifically, it’s proposed that when Glu_ext_ is protonated by the extracellular solution, both anion binding sites are occupied by F^-^. As Glu_ext_ rotates to face the intracellular solution and release its proton, the F^-^ in S_cen_ is also released intracellularly, and the now anionic sidechain of Glu_ext_ can occupy the vacant binding site. Glu_ext_ can then continue rotating up, pushing the other F^-^ into the extracellular solution, and a new F^-^ can enter S_cen_.^6^ Importantly, this proposed mechanism concludes a few points: (1) there should be anions occupying both binding sites at all times, and specifically there should be two F^-^ ions during key steps; (2) Glu_ext_ is able to directly interact with both the extra- and intra-cellular solutions to accept and release the transported proton; and (3) Glu_ext_ is performing the same role as E148 in the Cl^-^-transporting CLCs.

Interestingly, while there is no glutamate in an equivalent position as E203 in the Cl^-^ transporter, there is an additional glutamate (E318, hereafter referred to as Glu_int_) near Tyrcen and Metcen (Figure 1E). This glutamate has not been explored experimentally, but its existence has piqued the interest of researchers, who have begun trying to determine its role computationally. In 2021, Chon *et al*.^4^ performed a series of calculations on the crystal structures of wild type CLC^F^ and its mutants which determined Glu_int_ should remain deprotonated during the transport process and agreed with Last et *al*^6^ that Glu_ext_ is directly interacting with the extracellular and intracellular solutions. Later that same year, however, Chiariello et *al*^5^ concluded, through QM/MM simulations, that Glu_int_ is a part of the proton transport, but that while F^-^ is exported in its anionic form, the proton is released to the intracellular solution in the form of HF.

Because of these differences, we sought to determine new details about how this protein performs its anion export activity, and what the protonation states of Glu_ext_ and Glu_int_ should be to allow for the transport. Through the use of all-atom molecular dynamics (AA-MD) simulations and umbrella sampling calculations, we have determined a possible mechanism for the F^-^/H^+^ antiporting activity which can account for the previously seen experimental observations.

## Methods

### System Preparation

The protein structure was obtained from the RCSB Protein Data Bank (PDB ID: 6D0J^6^), and edited to its monomeric form. To determine the protonation state of all titratable residues at a pH of 7.4, with the exception of Glu_ext_ and Glu_int_, the H++ webserver^19–21^ was used. The protein was then embedded into a POPC bilayer and solvated with TIP3P water using Packmol-memgen^22^ and tleap in the Amber package^23^ to add the missing H atoms and NaCl ions to achieve a salt concentration of 0.150 M, resulting in systems of ~85,000 atoms. At this point, four systems were created, varying the protonation state of Glu_ext_ and Glu_int_: (1) both Glu_ext_ and Glu_int_ deprotonated; (2) only Glu_ext_ protonated; (3) only Glu_int_ protonated; (4) both Glu_ext_ and Glu_int_ protonated. Finally, tleap^23^ was used to generate the initial parameter and coordinate files for each of the four systems, using Amber’s ff14SB,^24^ lipid17,^25^ and TIP3P^26,27^ force fields, the latter of which also includes the atomic ions parameters used for fluoride.

### Molecular Dynamics Simulations

All simulations were conducted using the pmemd.cuda implementation of the Amber20 software package.^23,28,29^

In order to ensure proper relaxation and minimization of the structures, a multistep minimization procedure was used, each of which comprised of 5000 steps of steepest descent followed by 5000 steps of conjugate gradient minimization. First, only the water atoms were relaxed, with a 20 kcal/mol/Å^2^ restraint placed on all other atoms. Second, the ions and protein sidechains were also allowed to relax, with a 20 kcal/mol/Å^2^ restraint placed on only the protein backbone. Lastly, the entire system was allowed to relax, by removing all restraints.

After minimization, the systems were heated in two steps, each of which was 10 ps. First, they were heated from 0 K to 100 K, using the Langevin thermostat,^30^ during which time the protein, lipid, and F^-^ ions were held with a 10 kcal/mol/Å^2^ restraint. Second, they were heated from 100 K to 303 K, where, in addition to the Langevin thermostat, a Berendsen weak-coupling barostat^31^ was used to equilibrate the pressure, and again a 10 kcal/mol/Å^2^ restraint was placed on the protein, lipids, and F^-^ ions.

Next, ten equilibration steps were performed, each for 0.5 ns, to equilibrate the system’s periodic boundary condition dimensions. Finally, 200 ns of production simulations were run for each of the four systems. A time step of 2 fs was used, and all bonds involving hydrogen were constrained using the SHAKE algorithm. ^32^ The cutoff distance for long-range interactions was set to 12Å. The simulations were performed in an isothermal and isobaric ensemble, with the temperature held at 303 K, using the Langevin thermostat^30^ and Berendsen barostat. ^31^

### Umbrella Sampling

To determine the transport pathway used by F^-^ ions through the CLC^F^ protein, 200 snapshots over 200 ns from each simulation were analyzed using Caver 3.0.^33^ The lowest cost tunnel clusters were used to determine coordinates for F^-^ transport. For each of the four systems, umbrella sampling simulations were run with F^-^ ions spaced 0.3 Å apart through the tunnel, and held in place with a 30 kcal/mol/AÅ^2^ harmonic restraint. Each window ran for 10 ns, and the resultant trajectories were analyzed using the weighted histogram analysis method (WHAM).^34^

Umbrella sampling simulations were also run replacing F^-^ with HF for the system in which Glu_ext_ is deprotonated and Glu_int_ is protonated. The tunnel and window spacing used were the same as for the F^-^ simulations. The force field parameters for HF used were previously reported by Laage *et al*.^35^ and used for similar calculations by Yue *et al*.^36^

### Additional Analysis

All additional analyses were performed using the cpptraj program from the Amber20 software package.^37^ Root mean square deviation (RMSD) was calculated for the backbone atoms. The number of hydrogen bonds (*N_HBonds_*) were calculated, limited to only those found between the F^-^ ion and atoms in the CLC^F^ protein. Hydrogen bonds were also calculated between individual residues and the F^-^ ion during the umbrella sampling simulations, using only the windows when such interactions were possible (when the anion was within 3.3 Å of the donor atom(s) of the residue of interest). Hydration numbers (*N_Hydration_*) were calculated, and were defined as the number of water molecules within 3.5 A of the F^-^ ion which corresponds to its first solvation shell.^38^ The interaction energies were calculated and reported as the sum of the electrostatic and Van der Waals energies.

All visualization was performed using VMD software.^39^

## Results and Discussion

Initially, four AA-MD simulations were run using Amber20, varying the protonation state of Glu_ext_ and Glu_int_: (1) both Glu_ext_ and Glu_int_ deprotonated; (2) only Glu_ext_ protonated; (3) only Glu_int_ protonated; and (4) both Glu_ext_ and Glu_int_ protonated. All four systems behave similarly and quickly equilibrate, as determined by the RMSD shown in Figure S1. Interestingly, while the crystal structure of CLC^F^ indicates two binding sites, and as such we began our simulations with two F^-^ ions (F^-^_ext_ and F^-^_cen_), both sites do not remain occupied through the course of the simulation. In all four, as shown in the snapshots in Figures 2 and S2, F^-^_cen_ leaves the protein almost immediately. For the two systems in which Glu_ext_ is deprotonated, and therefore carries a negative charge, the remaining F^-^_ext_ ion is also pushed towards the intracellular side as Glu_ext_ rotates down to occupy the external binding site, in agreement with the windmill model proposed by Last *et al*.^6^ This rotation is shown in the snapshots, but is also highlighted in the dihedral angles of Glu_ext_ in Figure S3 which shows the system in which Glu_ext_ is protonated have constant dihedral angles, while those in which it is deprotonated have clear fluctuations. This same motion is tracked in Figure S4 by monitoring the hydrogen bonds between each of the F^-^ ions and the protein. This shows the rapid release of F^-^_cen_ (dashed lines) in all four systems, and the push of F^-^_ext_ (solid lines) in the deprotonated Glu_ext_ systems (black and teal lines). These results led to the conclusion that only a single F^-^ should be present in the binding area at a time, which can also help explain the 1:1 transport stoichiometry.

**Figure 2:**
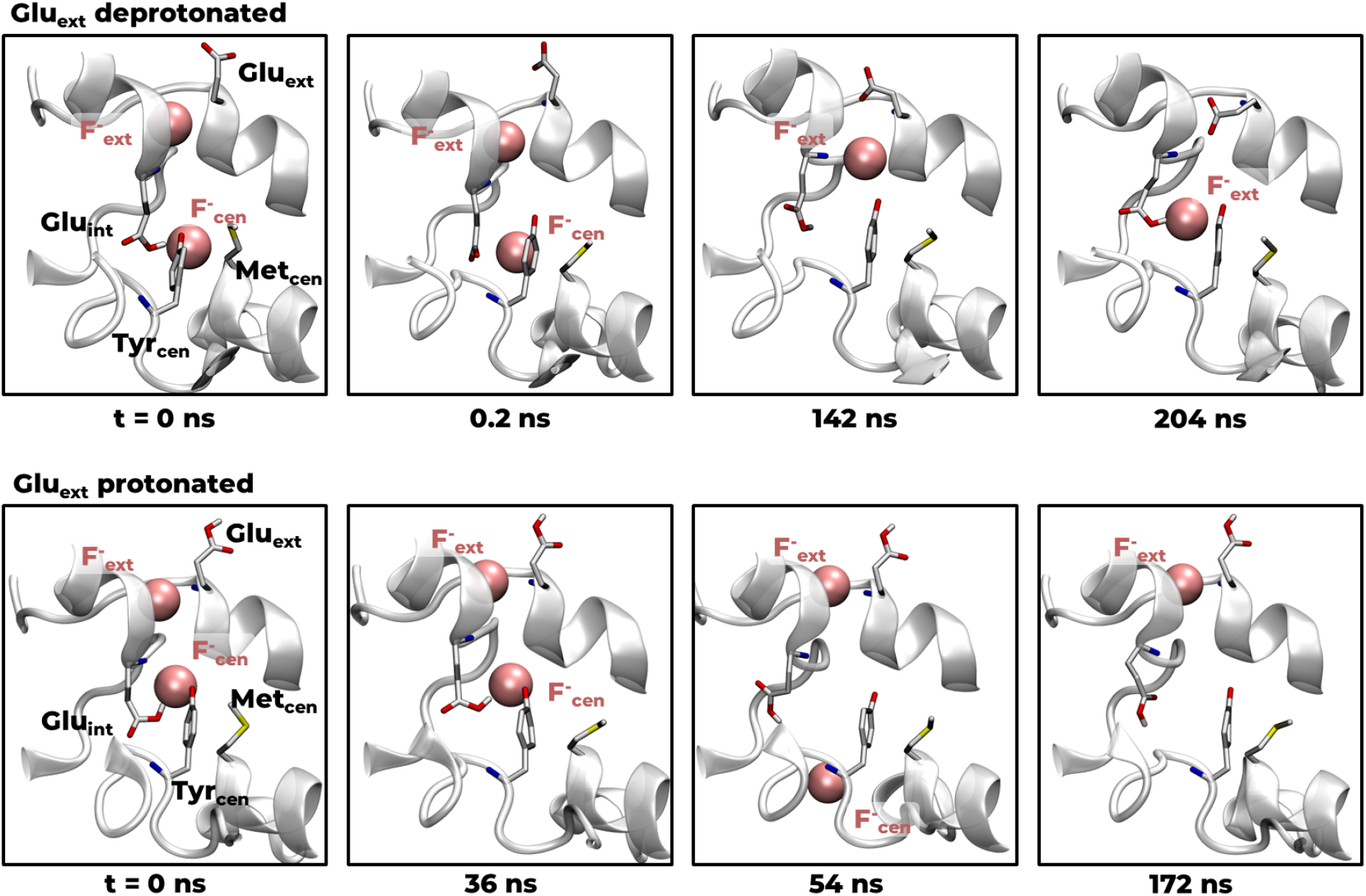
Anion movements when Glu_ext_ is deprotonated vs when it is protonated. For both series, snapshots were taken from the systems in which Glu_int_ is protonated, and snapshots for the other two systems are presented in Figure S2.

After these initial results, we sought to determine a comprehensive picture of the transport mechanism, and specifically how the proton import could be coupled to the F^-^ export by performing a series of umbrella sampling calculations to get the energetics of the F^-^ movement through the transport pathway. However, this pathway has not yet been well defined, therefore, the Caver Analyst software^33^ was first used to find this path. Using a single snapshot for each nanosecond of the production simulations to run through Caver, and comparing the ten lowest-cost tunnel clusters for each of the four systems, gives the results shown in Figures 3 and S5. In all four, the same pathway appears which encompasses the two known anion binding sites, and which can be used as the path for the umbrella sampling calculations. Interestingly, in the systems in which both Glu_ext_ and Glu_int_ have the same protonation state, there is a noticeable break in the pathway. In Figure S5A, when both are deprotonated, there is an empty space in the region connecting S_ext_ and S_cen_. In Figure S5C, when both are protonated, the pathway curves away from this same area. This gives the indication that these two states are not favorable for moving F^-^ through this central region of the transport pathway, but further analysis is necessary to make any conclusions.

**Figure 3:**
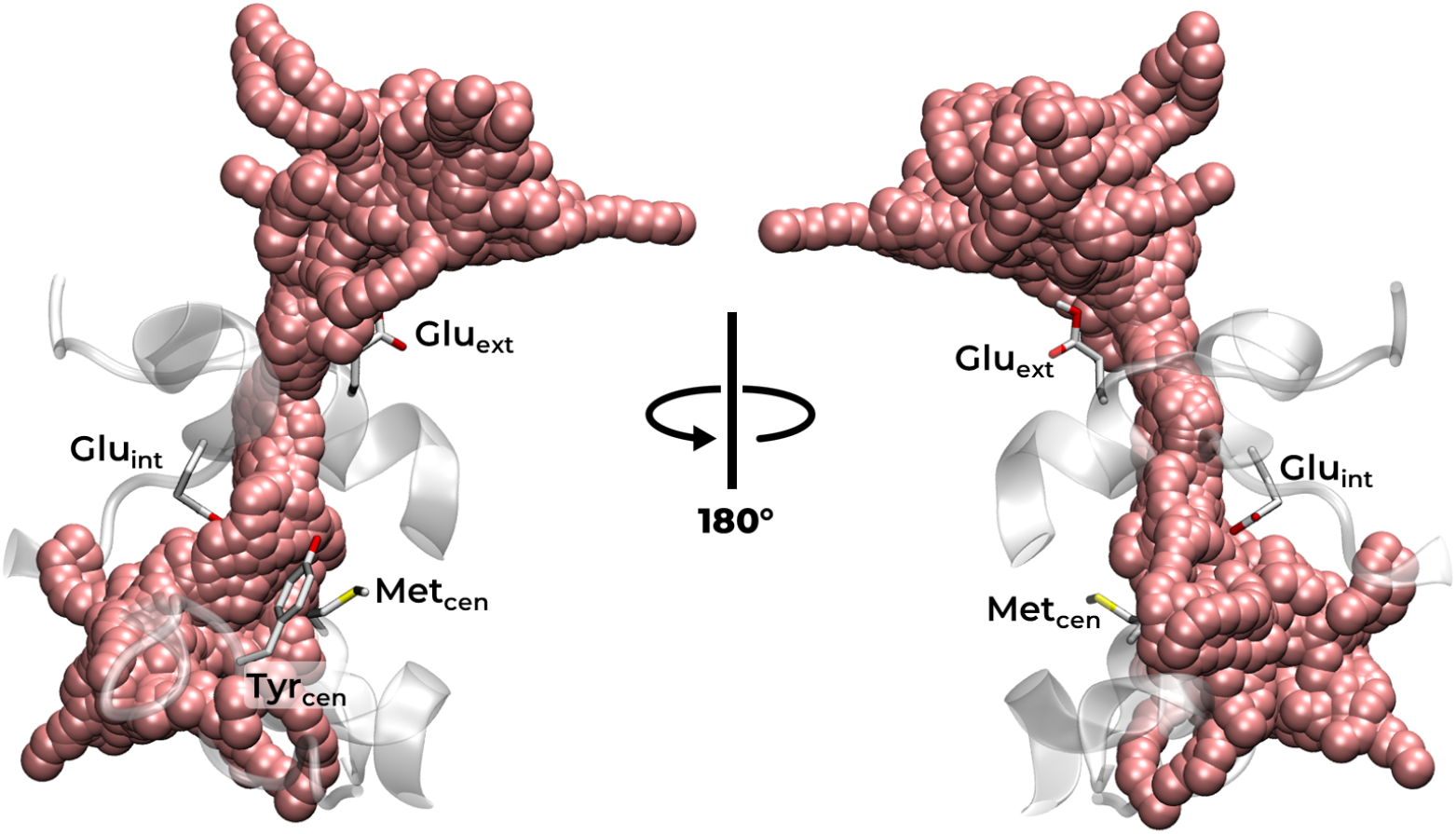
Top 10 tunnel clusters as determined by Caver analyst for the 200 ns trajectory of CLC^F^ in which Glu_ext_ is protonated and Glu_int_ is deprotonated, showing the pathway which encompasses the anion binding sites previously determined. The clusters for the other systems are presented in Figure S5.

Using the Caver results, we created 47 windows along the shared pathway, each with a single F^-^ ion, spaced ~0.3 A apart and held in place by a 30 kcal/mol/Å^2^ harmonic restraint. Separate calculations were performed for each of the four systems studied thus far, and the PMF of each is shown in Figure 4A as a function of the distance between F^-^ and the membrane center (ΔZ_COM_). Figure 4B shows the selection of the first and last windows (at ΔZ_COM_=-10.4 Å and ΔZ_COM_=3.5 Å, respectively) as beyond this point the ion is in direct contact with solution and any energies calculated by umbrella sampling would be inaccurate.

**Figure 4:**
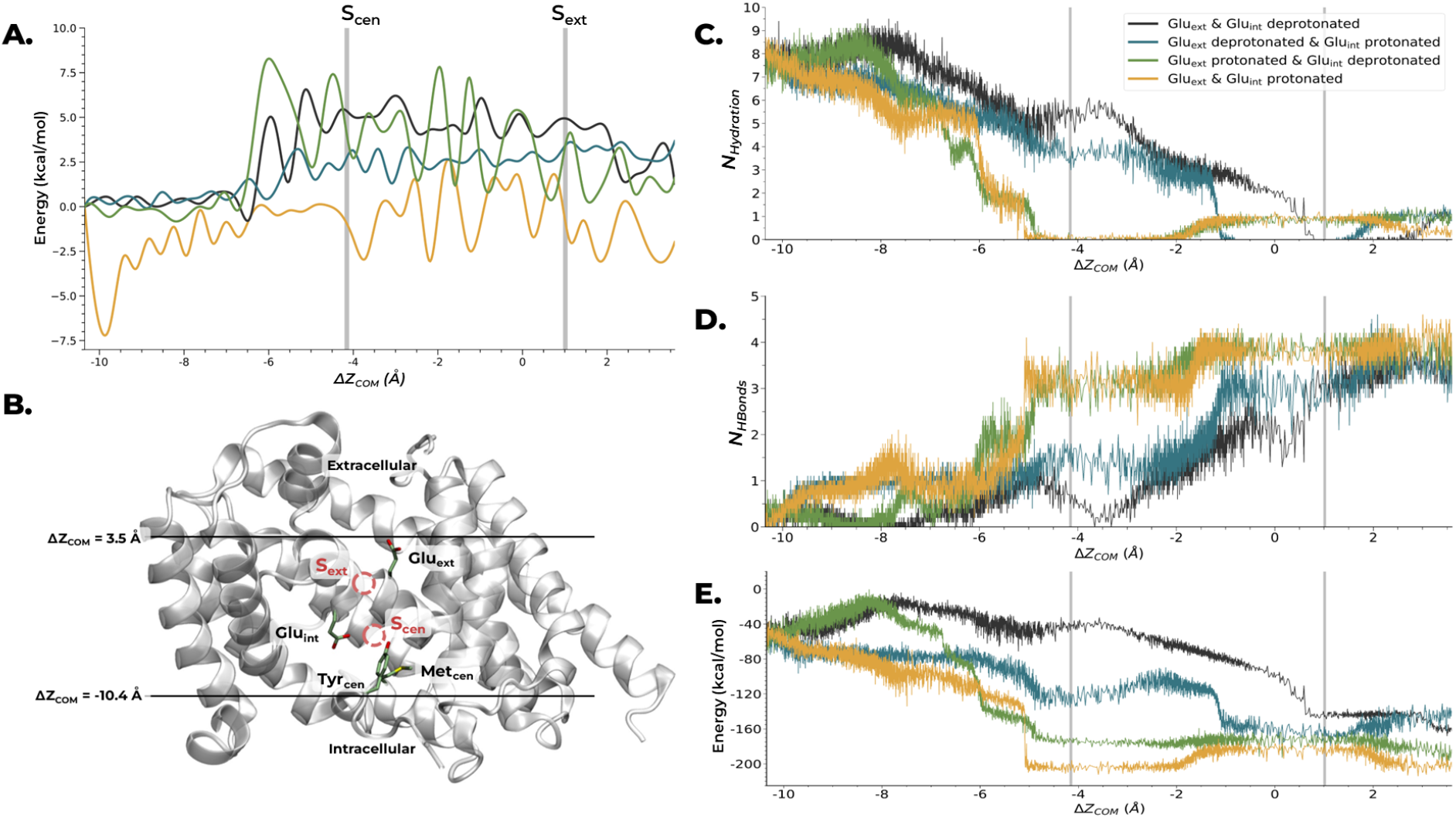
Results of umbrella sampling calculations performed on F^-^ transport through CLC^F^. A. PMF for all four systems, gray lines highlight the binding sites as determined by the crystal structure; B. Diagram of CLC^F^ protein, showing the bounds used for the umbrella sampling windows and the binding sites; C. Hydration number for F^-^ in each of the four systems; D. Number of hydrogen bonds between F^-^ and CLC^F^ protein in each of the four systems; E. Total interaction energy (sum of electrostatic and van der Waals energies) between F^-^ and CLC^F^ protein for each of the four systems.

These calculated PMFs reveal a number of details about the transport mechanism of CLC^F^. Most noticeably, the yellow curve, when both Glu_ext_ and Glu_int_ are protonated, seems to be overstabilized throughout the course of the PMF. At multiple points through the transport pathway, this curve has energy barriers of 3-6 kcal/mol in order to move on, which is unfavorable, especially when compared to the much lower energy barriers calculated in the other systems. Additionally, the green curve has a slightly lower energy (−0.1 kcal/mol compared to 0.4 and 0.5 kcal/mol for the black and teal curves, respectively) at the beginning of the PMF. This indicates an initial preference for Glu_ext_ to be protonated and Glu_int_ to be deprotonated as F^-^ enters the pathway. The green curve remains at a lower energy until ΔZ_COM_=-6.5 A, at which point it spikes to 7.8 kcal/mol. Comparing the other systems, the teal curve, when only Gluint is protonated, gives the lowest energy barriers (0-2 kcal/mol) for the remainder of the PMF. In the last few windows, the black curve representing the doubly deprotonated system has a drop in energy, however as the umbrella sampling simulations couldn’t be continued past this point, it’s difficult to make a definite conclusion.

These results allow us to piece together a proton transfer mechanism that allows for the minimization of energy barriers. For F^-^ to enter the pathway, Gluext should be protonated, while Glu_int_ is deprotonated. When ΔZ_COM_=-6.5 Å, shortly before F^-^ enters S_cen_, a proton transfer can occur, making Gluext deprotonated and Gluint protonated, allowing for energy minimization. This protonation state remains the favorable state, until ΔZ_COM_ ≈3 A, at which point it’s possible the system where both Glu_ext_ and Glu_int_ are deprotonated is more favorable.

While these PMFs provide significant insight on their own, a more detailed understanding of the mechanism requires further analysis. We determined the hydration number (*N_hydration_*) as the number of water molecules within 3.5 A of F^-^ over the course of all the umbrella sampling simulations in Figure 4C. Upon initial entrance of F^-^ into the tunnel, all of the systems begin with 8 water molecules coordinating the anion. From this entrance until ΔZ_COM_=-8.2 A, the green and black curves both show a stable, if not slightly increasing, *N_hydration_* while the yellow and teal curves show a slight dehydration as *N_hydration_* drops from 8 to 5-6 water molecules. From ΔZ_COM_=-8.2 Å until ΔZ_COM_=-5 Å, all four systems become dehydrated, however the yellow and green curves have a much steeper slope than the black and teal curves, the latter of which don’t show complete dehydration until ΔZ_COM_ is 1 A and −1 Å, respectively. It’s important to note that fluoride is known to have a very large hydration energy (≈ 111 kcal/mol^40^), and therefore rapid and complete dehydration is unfavorable, unless there is a compensatory stabilization.

To determine how F^-^ is overcoming this dehydration, we also measured the number of hydrogen bonds in Figure 4D between the ion and CLC^F^. The results show all four curves have mirrored slopes compared to the *NH_ydration_* curves in Figure 4C, indicating that as F^-^ is losing its coordinating water, it is making up for it by forming hydrogen bonds with residues from CLC^F^. Knowing that F^-^ is forming these interactions with the protein, the next question is if these interactions are strong enough to overcome the hydration energy. In Figure 4E, we calculated the total interaction energy between F^-^ and the protein, and the curves match the curve trends observed in Figure 4C. Again, the green and black curves stay more stable until ΔZ_COM_=-8.2 Å, while the yellow and teal curves immediately begin to decrease, from −40 kcal/mol to −100 kcal/mol. At this point, until ΔZ_COM_ ≈-5 A, all four curves decrease, again with the yellow and green showing a steeper slope. From ΔZ_COM_=-5 A to ΔZ_COM_=-2 A the yellow curve stays constant at a lower value (−200 kcal/mol) than the others, but then has an increase in energy bringing it to the same energy as both the teal and green curves around −160 kcal/mol. Once ΔZ_COM_ ≈2-2.5 A all of the curves show a slight energy decrease, except for the teal curve which shows a slight increase.

Connecting these analyses with the PMF, the protonation states of and proton transfer between Gluext and Gluint can be determined. Based on the overstabilization of the doubly protonated state (yellow curve in Figure 4A) and its consistently large energy barriers, its rapid dehydration, and the previous Caver results which also disfavor this system, it seems very unlikely for this state to exist during the transport process. Examining the other three systems, then, the PMF indicates an initial preference for Gluext to be protonated and Gluint to be deprotonated (green curve in Figure 4A). This remains the more favorable state, until shortly before F^-^ reaches S_cen_, at which point a proton transfer from Glu_ext_ to Glu_int_ (teal curve in Figure 4A) would avoid the large energy barrier of 7.5 kcal/mol at ΔZ_COM_=-6.5 A. At this same point, the system in which both Gluext and Gluint are deprotonated (black curve in Figure 4A) has an energy barrier of 6.0 kcal/mol, which indicates only the system in which Glu_int_ is protonated (teal curve) has a reasonably low barrier to overcome. The hydration data for the corresponding region indicates the system with only Glu_ext_ protonated (green curve in Figure 4C) maintains its hydration for longer, but when ΔZ_COM_=-6.5 A, there is a rapid dehydration for this system, whereas a proton transfer from Glu_ext_ to Glu_int_ (teal curve in Figure 4C) allows for a much slower dehydration. Because of fluoride’s high hydration energy, this could explain why the PMF showed a significantly lower energy barrier for the teal curve compared to the green curve (1 kcal/mol versus 7.5 kcal/mol) at this point. From ΔZ_COM_=-6.5Å until ΔZ_COM_ ≈3 Å, the PMF shows this system in which Glu_ext_ is deprotonated and Gluint is protonated (teal curve in Figure 4A) remains the one with the lowest energy barriers. From ΔZ_COM_ ≈3 Å until the end of the explored pathway at ΔZ_COM_=3.5 Å, there is a stabilization of the doubly deprotonated system (black curve in Figure 4A), which matches the beginning of F^-^ becoming rehydrated (Figure 4C). As stated previously, beyond this point the extracellular solution is in direct contact with F^-^ and the umbrella sampling simulations couldn’t continue without seeing unrealistically large energy barrier.

Next, we wanted to get an understanding of the role played by individual residues in this transport mechanism. In order to identify which residues were playing a role, the frequency of hydrogen bond interactions between the F^-^ ion and each residue were quantified in Figure S6. Using these and the previous results as a guide, we visualized the simulations and determined the mechanism of transport shown in Figure 5.

**Figure 5:**
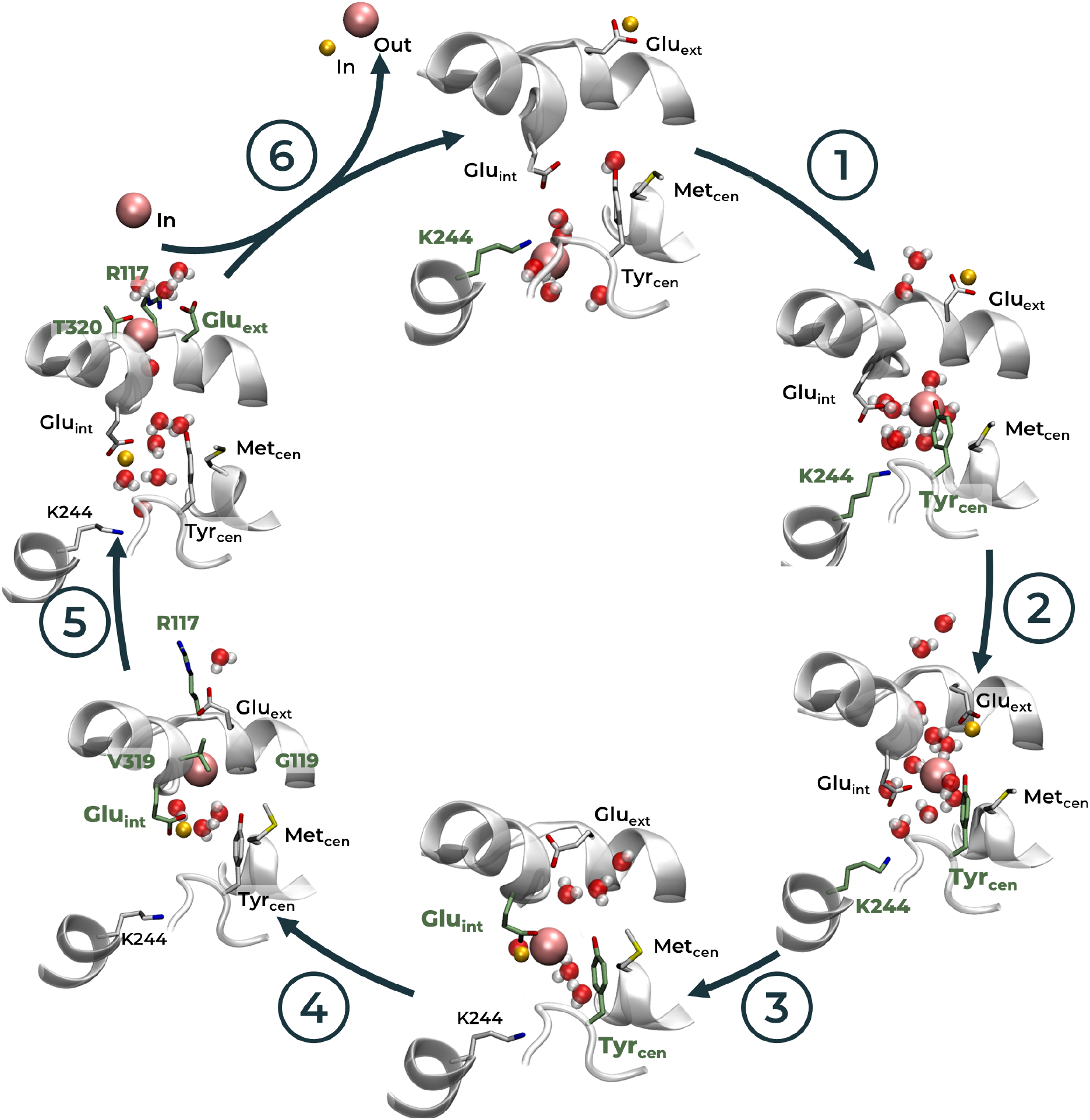
Transport mechanism of CLC^F^ F^-^/H^+^ antiporter. Residues coordinating F^-^ at each step shown in green licorice, and other important residues shown in white licorice. Only water molecules within 5 Å of F^-^, Glu_ext_, or Glu_int_ are shown.

To recruit F^-^ into the transport pathway, the sidechain of Lys244 coordinates the anion and pulls it up towards Tyr_cen_ (Figure 5–1), which can then coordinate it. The F^-^ remains coordinated by several waters, though some are being replaced by these coordinating residues, matching the *Nhydration* and *N_HBond_s* data in Figures 4C and 4D. At this point, F^-^ is almost to the central binding site, but before it reaches there is a rotation of Glu_ext_ down to face the anion (Figure 5–2). This rotation of Glu_ext_ is tracked in Figure S7, where the green curve fluctuates between facing the extracellular solution and stretching down towards S_cen_. In this position, water flows into the binding area, where it can facilitate the proton transfer between Glu_ext_ and Glu_int_ (Figure 5–3). Next, F^-^ briefly sits in the central binding site, coordinated by Tyrcen and the now protonated Gluint, before moving up to occupy the external binding site (Figure 5–4). As it moves up, the repulsion between itself and the now negatively-charged Gluext forces the residue to continue rotating until it is again facing the extracellular solution. To overcome this repulsion, and allow the F^-^ to continue moving instead of being pushed back down, several residues begin coordinating it, including the sidechains of Thr320, Arg117, and the backbones of Gly119, Val319, and Glu_ext_ (Figure 5–5) which matches the hydration and hydrogen bonding data of Figures 4C and 4D where the F^-^ becomes fully dehydrated and is coordinated by 4-5 residues (teal curve, ΔZ_COM_=-1 A). Once F^-^ is in the external binding site and Glu_ext_ is facing up and out, water can easily access F^-^, rehydrating it as it joins the extracellular solution (Figure 5–6). At the same time, Glu_int_ is in direct contact with the intracellular solution, which could allow it to release the proton and complete the antiporter activity, an observation we cannot confirm as it is beyond the boundaries of the umbrella sampling simulations.

This mechanism left a few questions unanswered, however. What is the role of Metcen if it isn’t directly interacting with F^-^, and why is a negatively charged Gluint favorable when the anionic F^-^ is entering the channel near it? To answer both, we looked at the interactions both of these residues are forming throughout the simulations, the results of which are in Figures S8 and S9. In Figure S8, the interaction between Met_cen_ and Tyr_cen_ is tracked, as the interaction between methionine and aromatic residues has been well-documented in stabilizing proteins.^41^ Figure S8 shows that these residues remain in contact (defined as a sulfur-aromatic ring distance ¡7 Å^41^) through the duration of all of the simulations, which, in turn, keeps Tyr_cen_ in a stable conformation, as tracked by its χ_1_ dihedral angle. Additionally, when tracking the hydrogen bonds formed by Gluint in Figure S9, while it interacts with Leu322 consistently which maintains its alpha helical structure, when it is deprotonated it is able to form hydrogen bonds with both Tyr_cen_ and Lys244. The interaction with Lys244 is also shown in Figure S10 when tracking the distance between Glu_int_ and Lys244. When F^-^ is initially entering the binding area, and when the proposed mechanism indicates Lys244 should be forming a hydrogen bond with the anion, Lys244 is also interacting with the deprotonated Glu_int_, shown by a decrease in the distance of 1.4 and 1.1 Å between the two residues in the black and green curves, respectively (Figure S10). At the same time, there is a decrease in the distance between Lys244 and F^-^ of 1.0 and 0.8 ^Å^ in the same two systems, indicating that Lys244 is better able to interact with F^-^ when it is being held by Gluint. The interaction between Glu_int_ and Tyr_cen_ was similarly tracked in Figure S11. As the Glu_int_ - Lys244 interaction is ending (ΔZ_COM_=-9 Å in Figure S10), there is a decrease in the distance of 0.7 and 0.5 Å between Glu_int_ and Tyr_cen_ for the systems in which Glu_int_ is deprotonated (black and green curves, respectively, in Figure S11). Because Tyr_cen_ is being held rather rigidly by Met_cen_, this indicates Glu_int_ is moving away from K244 and towards Tyr_cen_. As it does so, the F^-^ ion briefly moves further away from Tyr_cen_ by 0.6 and 0.5 A, likely propelled by the electrostatic repulsion between it and Glu_int_. This motion can help further explain why F^-^ would continue to move into the transport pathway after its brief interaction with Lys244, instead of rejoining the intracellular solution.

Finally, we sought to ensure fluoride was being transported in its anionic form and not as the more common HF. Therefore, we ran the same umbrella sampling simulations replacing the anion with HF, and calculated the corresponding PMF. Specifically, we performed the simulations on the system in which Glu_ext_ is deprotonated and Glu_int_ is protonated as it consistently had very low energy barriers for F^-^ transport. The results in Figure S12 show that while the range of energies for the system containing F^-^ was −0.1 to 3.6 kcal/mol (teal curve in Figure S12), for the same system containing HF, the range jumps to −7.7 to 32.9 kcal/mol (pale blue curve in Figure S12). This shows that HF transport through CLC^F^ is highly improbable, as these energy barriers are incredibly large.

## Conclusions

In summary, we have used MD simulations combined with umbrella sampling simulations to determine a reasonable mechanism for the proton-coupled transport of F^-^ in the CLC^F^ antiporter protein. Our simulations reveal that only a single F^-^ anion should be present in the protein’s transport pathway at a time, which helps explain the unique 1:1 anion:proton stoichiometry that sets CLC^F^ apart from the other CLC antiporters. Our umbrella sampling results also reveal that Glu_ext_ (Glu118) maintains the same rotational gating function as is shown in Glu148 in the CLCs, and which corresponds with the windmill model proposed by Last *et al*.^6^ Our proposed mechanism shows the importance of several residues which were previously reported to be necessary for F^-^ efflux, experimentally, including Val319, Thr320, and Tyr396,^6^ and, most notably, indicates Glu_int_ (Glu318) is able to receive the proton from Gluext before passing it on to the intracellular solution. The PMFs calculated here result in energy barriers of 0.1-2.6 kcal/mol, which is comparable to the PMFs previously reported for the CLC Cl^-^/H^+^ antiporter which gave energy barriers of between 2 and 4 kcal/mol. ^42,43^ Additionally, we have shown that CLC^F^ is transporting fluoride as an anion and not as HF, as the latter creates very large energy barriers for the system to overcome, a result which complements the recent work by Yue *et al*.^36^ on the Fluc F^-^ channel.

This work is among the first reported computational studies of the CLC^F^ antiporter protein, and is the first to propose a mechanism to couple the proton import with F^-^ export. The previous computational studies were contradictory in their proposed role of Gluint, and our study has provided evidence that this residue is necessary for proton import as its changing protonation state allows for the minimization of energy barriers.

## Data Availability

Input files for the molecular dynamics simulations and umbrella sampling calculations, including parameter and initial coordinate files, are available at DOI: 10.5281/zenodo.7117547.

## Author Contributions

H.T. designed and supervised the project; K.M. performed the computational studies and analyses; H.T. and K.M. analyzed the data and wrote the manuscript together.

## Acknowledgements

The authors acknowledge the Texas Advanced Computing Center (TACC) at The University of Texas at Austin as well as the Office of Information Technology Cyberinfrastructure Research Computing (CIRC) at The University of Texas at Dallas for providing HPC resources that have contributed to the research results reported within this paper.

