## Supplemental Information for "Uncovering the Mechanism of the Proton-Coupled Fluoride Transport in the CLC^F^ Antiporter"

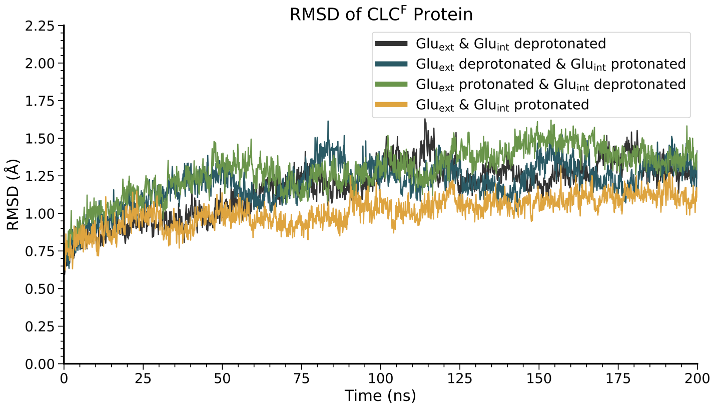


Figure S1: RMSD for initial simulations on all four systems of CLC$^F$

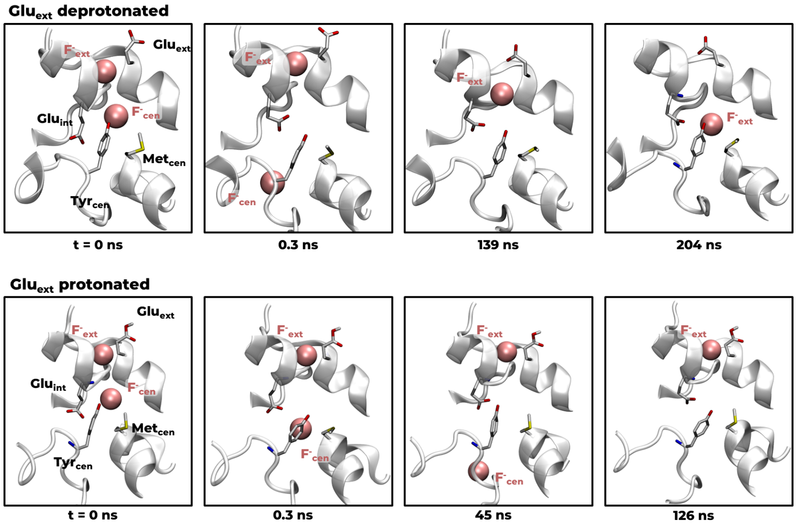


Figure S2: Snapshots showing ion movement in the two CLC$^F$ MD simulations in which Glu$_{int}$ is deprotonated.

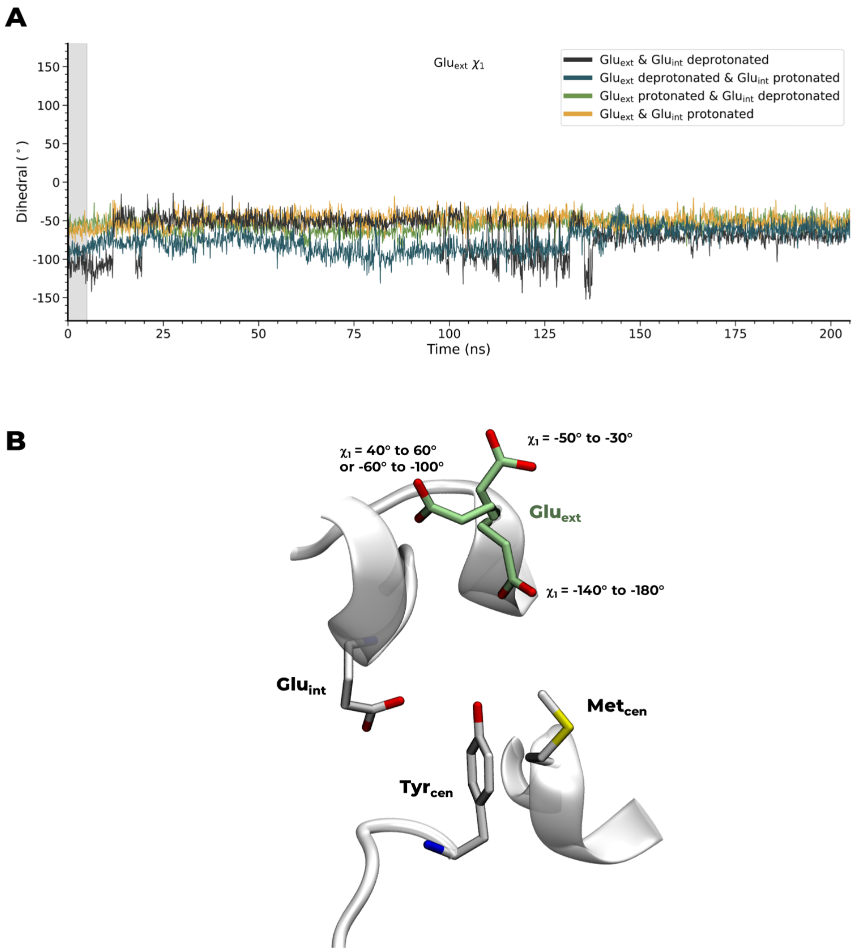


Figure S3: A. Dihedral angles of Glu$_{\text{ext}}$ during CLC$^{\text{F}}$ MD simulations. The gray shaded region indicates the equilibration steps of the simulation. B. Representation of Glu$_{\text{ext}}$ orientation at different dihedral angles during the simulations. Glu$_{\text{ext}}$ is highlighted in green.

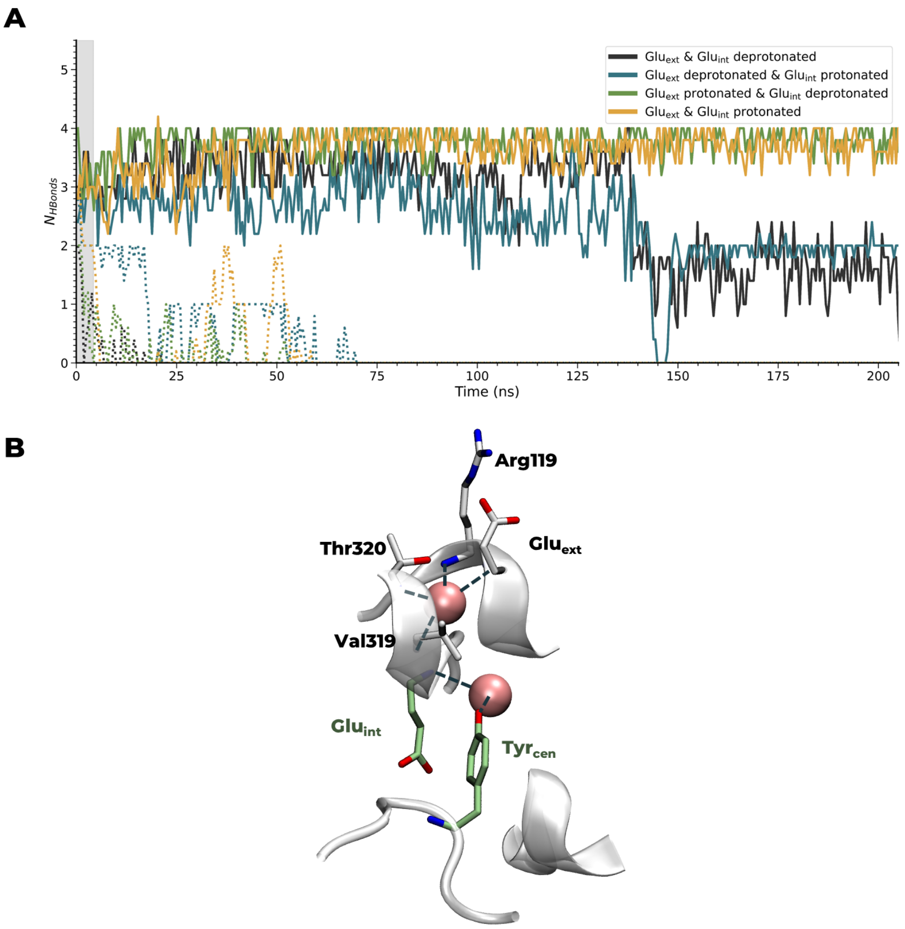


Figure S4: A. Number of hydrogen bonds between the $F^-$ ions and $CLC^F$ protein. Solid lines represent $F^-{}_{ext}$ and dashed lines represent $F^-{}_{cen}$. The gray shaded region indicates the equilibration steps of the simulation. B. Snapshot showing the hydrogen bonds between $F^-$ and $CLC^F$ in each of the binding sites. Residues involved in $S_{ext}$ are shown in white and residues involved in $S_{cen}$ are shown in green.

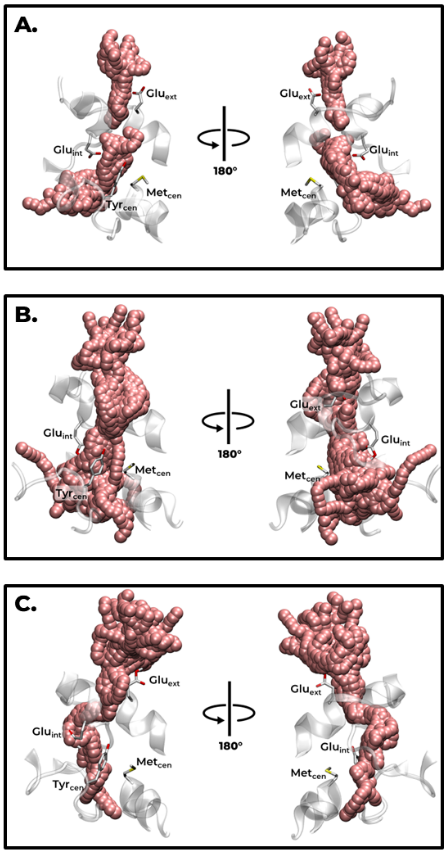


Figure S5: Top 10 tunnel clusters as determined by Caver analyst for each of the systems' trajectories, showing the same pathway in each which encompasses the anion binding sites previously determined. A. System when Glu$_{ext}$ and Glu$_{int}$ are deprotonated; B. System when Glu$_{ext}$ is deprotonated and Glu$_{int}$ is protonated; C. System when Glu$_{ext}$ and Glu$_{int}$ are protonated

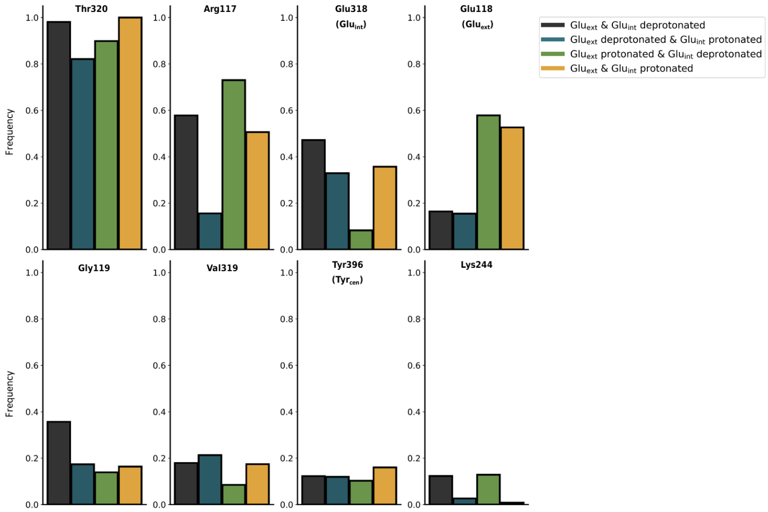


Figure S6: Frequency of hydrogen bonds between F$^-$ ion and individual residues during the umbrella sampling simulations.

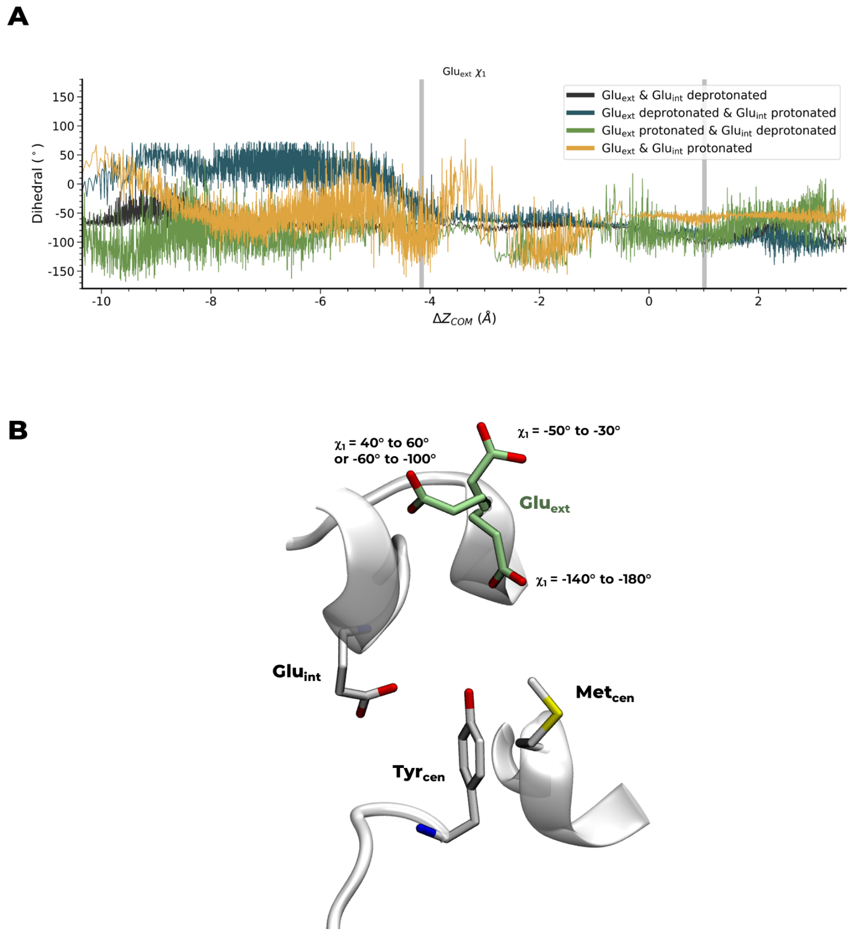


Figure S7: A. Dihedral angles of Glu$_{ext}$ during umbrella sampling calculations. The gray lines indicate the S$_{cen}$ and S$_{ext}$ binding sites. B. Representation of Glu$_{ext}$ orientation at different dihedral angles during the simulations. Glu$_{ext}$ is highlighted in green.

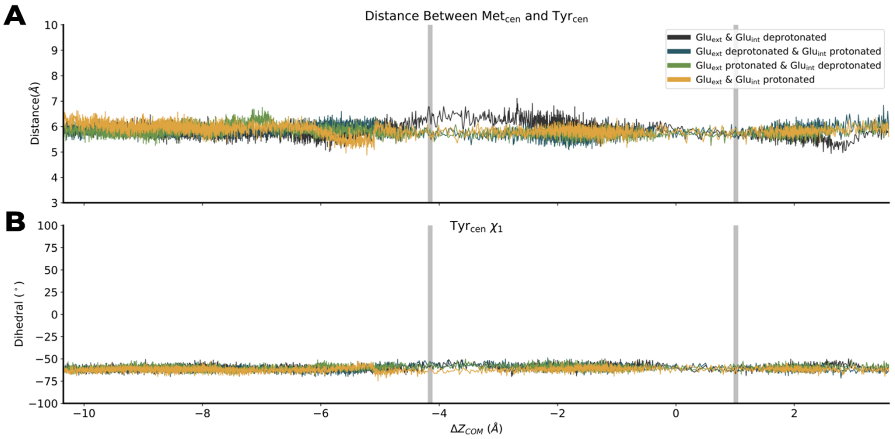


Figure S8: The relationship between Met$_{\mathrm{cen}}$ and Tyr$_{\mathrm{cen}}$. A. The distance between the sulfur of Met$_{\mathrm{cen}}$ and the center of mass of the aromatic ring in Tyr$_{\mathrm{cen}}$. B. The $\chi_1$ dihedral angle of Tyr$_{\mathrm{cen}}$.

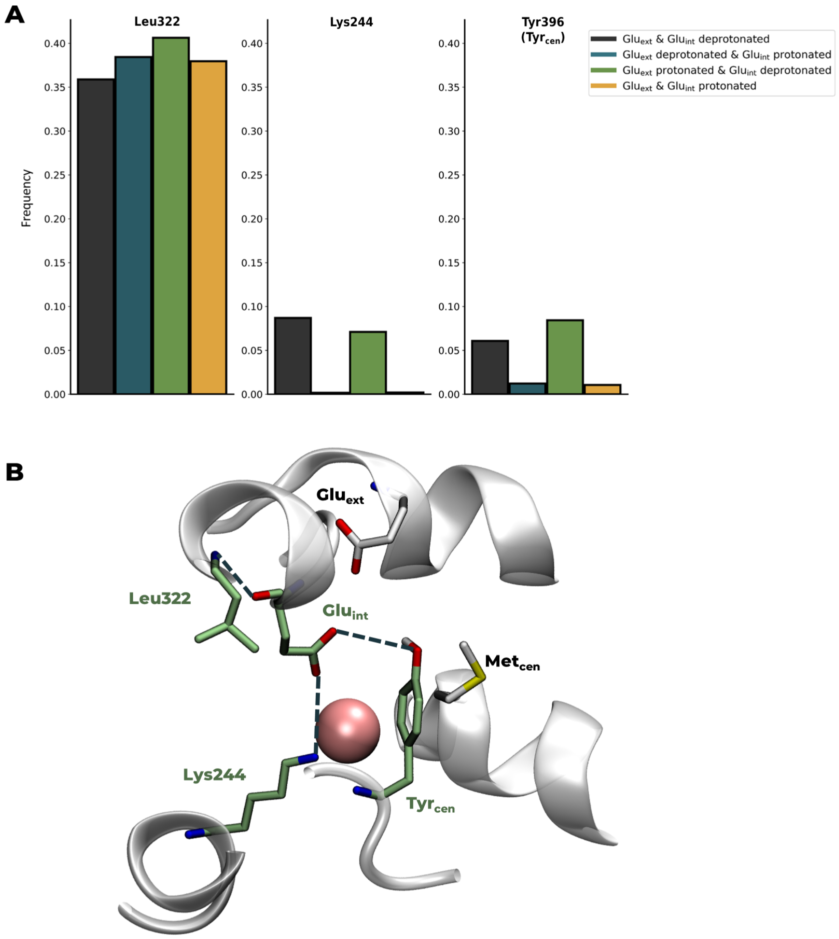


Figure S9: A.The frequency of hydrogen bonds between $\mathrm{Glu_{int}}$ and other $\mathrm{CLC^F}$ residues. B. A representation of the hydrogen bonds between $\mathrm{Glu_{int}}$ and other $\mathrm{CLC^F}$ residues

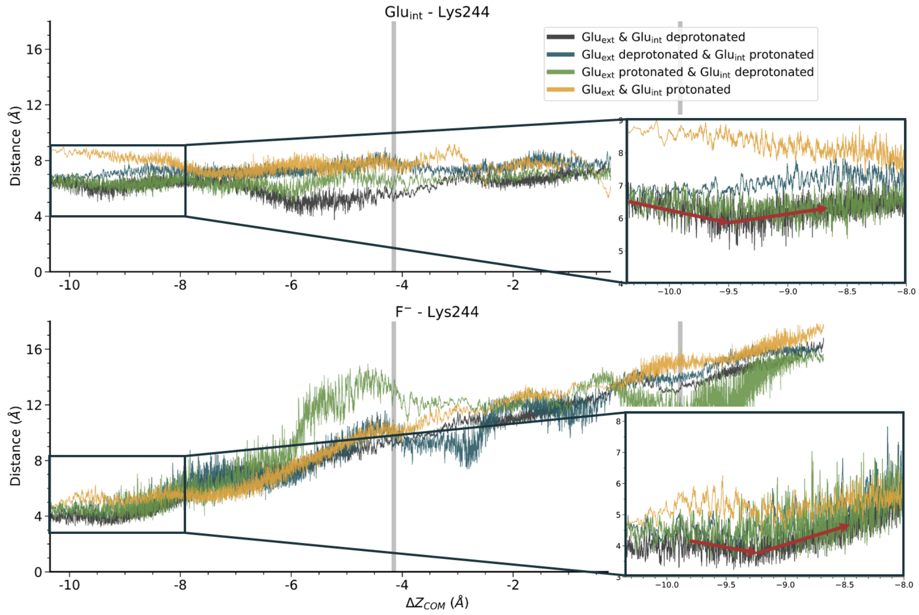


Figure S10: The distance between the sidechain oxygen atoms of Glu$_{\text{int}}$ and the sidechain nitrogen atom of Lys244 and the distance between the sidechain nitrogen of Lys244 and the F$^-$ ion. Highlighted in the zoomed-in region of the plot is the change in distance for the green and black curves. In the green curve, the distance between Glu$_{\text{int}}$ and Lys244 decreases from 7.1 to 5.5 Å from $\Delta Z_{\text{COM}}$=-10.35 to -9.5 Å before increasing back to 6.9 Å at $\Delta Z_{\text{COM}}$=-8.5 Å. During the same region of the plot, the black curve decreases from 6.6 to 5.5 Å before increasing to 6.9 Å. Similarly, for the distance between Lys244 and the F$^-$ ion, there is a decrease in the green curve from 4.5 to 3.4 Å followed by an increase to 4.7 Å. For the black curve, this distance decreases from 4.6 to 3.9 Å before increased to 5.3 Å.

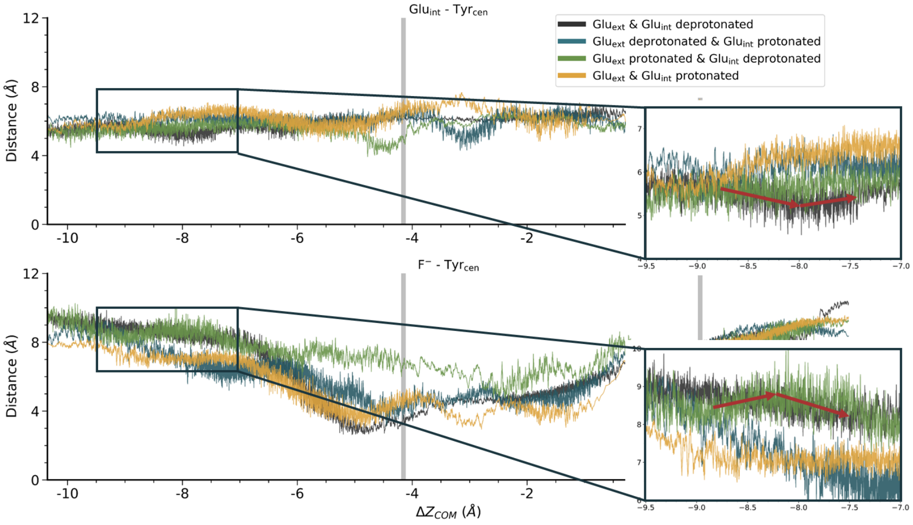


Figure S11: The distance between the sidechain oxygen atoms of $\text{Glu}_{\text{int}}$ and the sidechain oxygen atom of $\text{Tyr}_{\text{cen}}$ and the distance between the sidechain oxygen of $\text{Tyr}_{\text{cen}}$ and the $\text{F}^{-}$ ion. Highlighted in the zoomed-in region of the plot is the change in distance for the green and black curves. In the green curve, the distance between $\text{Glu}_{\text{int}}$ and $\text{Tyr}_{\text{cen}}$ decreases from 5.5 to 5.0 Å from $\Delta Z_{\text{COM}}$=-9 to -7.8 Å before increasing back to 5.7 Å at $\Delta Z_{\text{COM}}$=-7 Å. During the same region of the plot, the black curve decreases from 5.7 to 5.1 Å before increasing to 5.6 Å. For the distance between $\text{Tyr}_{\text{cen}}$ and the $\text{F}^{-}$ ion, there is an increase in the green curve from 8.2 to 9 Å followed by a decrease to 8.1 Å. For the black curve, this distance increases from 8.5 to 8.9 Å before decreasing to 8.1 Å.

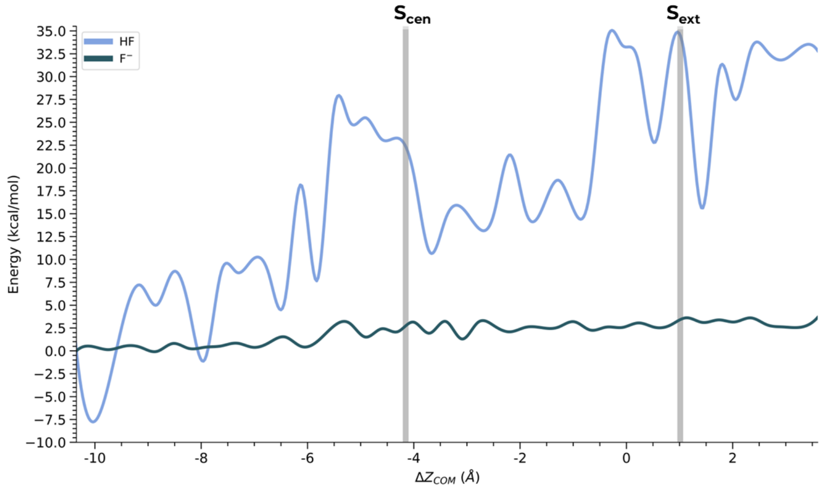


Figure S12: PMF comparison between the transport of $F^-$ ion and HF. For both systems, $Glu_{ext}$ is deprotonated and $Glu_{int}$ is protonated. Gray lines highlight the binding sites as determined by the crystal structure.
